# Functional microRNA targetome undergoes degeneration-induced shift in the retina

**DOI:** 10.1101/2020.05.27.118307

**Authors:** Joshua A. Chu-Tan, Zhi-Ping Feng, Yvette Wooff, Adrian V. Cioanca, Ulrike Schumann, Riemke Aggio-Bruce, Hardip Patel, Matt Rutar, Katherine Hannan, Konstantin Panov, Jan Provis, Riccardo Natoli

**Author notes:** Corresponding author: Riccardo Natoli, Clear Vision Research Laboratory, Eccles Institute of Neuroscience, John Curtin School of Medical Research, College of Health and Medicine, The Australian National University, Acton, ACT 2601, Australia.

## Abstract

MicroRNA (miRNA) play a significant role in the pathogenesis of complex neurodegenerative diseases, including age-related macular degeneration, acting as post-transcriptional gene suppressors through their association with argonaute (AGO) protein family members. However, to understand their role in disease, investigation into the regulatory nature of miRNA with their targets is required. To identify the active-miRnome-targetome interactions in the degenerating retina, AGO2 HITS-CLIP was performed using a mouse model of retinal degeneration. Analysis revealed a similar miRnome between healthy and damaged retinas, however, a shift in the active targetome was observed. This shift was also demonstrated by a change in the seed binding regions of miR-124-3p, the most abundant retinal miRNA loaded in AGO2. Following damage, AGO2 was localised to the inner retinal layers indicating a locational miRNA response to retinal damage. This study provides important insight into the alteration of miRNA regulatory activity that occurs as a response to retinal degeneration.

## Introduction

MicroRNA (miRNA) are a class of endogenous non-coding RNAs, implicated in the control of cellular and tissue homeostasis, development and biological pathway regulation (Krol et al., 2010), acting as post-transcriptional gene repressors. In order for miRNA to exert their biological effects they are complexed with Argonaute (AGO) proteins as part of a larger RNA-induced silencing complex (RISC) (Elbashir et al., 2001a, Elbashir et al., 2001b, Hammond et al., 2000, Martinez et al., 2002, Nykanen et al., 2001, Schwarz et al., 2002). Once incorporated into RISC, the miRNA canonical activity mediates binding to the 3’untranslated (3’UTR) of the mRNA target(s) through the recognition of the miRNA seed regions (typically 6-8 nucleotides long, commonly referred to as 6-mer, 7-mer and 8-mer seed regions) (Lewis et al., 2005). This process results in their functional capabilities of transcript silencing through translational repression or mRNA degradation (Bazzini et al., 2012, Djuranovic et al., 2012).

A single miRNA has the ability to target hundreds of mRNAs, often all working in similar biological pathways, thus allowing for the mapping and identification of specific “miRNA networks” (Friedman et al., 2009). Due to their selective targeting pattern and ability, miRNA have emerged as key orchestrators of the mammalian transcriptome and as such, their dysregulation has been implicated in the pathogenesis of multiple inflammatory diseases, cancers, neurological disorders, and retinal degenerative diseases including Age-Related Macular Degeneration (AMD) (Bartel, 2004, Bartel and Chen, 2004, Lukiw et al., 2012, Qiu et al., 2015, Zhou et al., 2014, Chu-Tan et al., 2018).

AMD is a degenerative disease of the central retina, and is the leading cause of blindness in the developed world (Ambati et al., 2003). Dry AMD, the more prevalent and currently untreatable form, is characterised by the focal death of the light-sensing photoreceptor cells, and underlying retinal pigmented epithelium (RPE) in the retina, which results in progressive and irreversible vision loss. Due to a complex and multi-faceted aetiology, causal links to AMD remain elusive (Ambati et al., 2003, Ambati et al., 2013, Beatty et al., 2000, Hollyfield et al., 2008, Hollyfield et al., 2010, Salomon et al., 2011). Whilst Genome-Wide Association Studies (GWAS) have identified major oxidative stress and inflammatory pathways as key processes involved in AMD disease onset and progression, these findings alone have not proved efficacious in regard to therapeutic development, because targeting single proteins does not mediate a holistic therapeutic for the disease ((Klein et al., 2005), reviewed in (Kassa et al., 2019, Park et al., 2019)). For these reasons, it may prove more valuable to invest in the therapeutic manipulation of multiple targets within known dysregulated pathways, drawing attention to the use of miRNA as potential therapeutic options for the treatment of AMD.

Multiple miRNAs have been postulated to play a role in the pathogenesis of AMD (Askou et al., 2018, Berber et al., 2017, Bhattacharjee et al., 2016, Ertekin et al., 2014, Grassmann et al., 2014, Lukiw et al., 2012, Menard et al., 2016, Murad et al., 2014, Szemraj et al., 2015, Zhou et al., 2014, Chu-Tan et al., 2018). One example is miR-124 which has been heavily studied in the central nervous system (CNS) and is the most highly abundant miRNA in the brain (Conaco et al., 2006, Lagos-Quintana et al., 2002, Lim et al., 2005, Makeyev et al., 2007). However, there is inconsistency across publications in terms of elucidating functionally relevant miRNAs and their targets, largely due to an overprediction of possible miRNA binding sites. It is therefore imperative that the miRNA transcriptome (miRnome) and mRNA targetome are elucidated, in both the healthy and diseased retina, in order to fully understand their role and potential in the therapeutic space.

In order to further advance our understanding of the molecular roles that miRNA, including miR-124, play in the normal and degenerating retina, we used a modified version of the HITS-CLIP (High-Throughput Sequencing following Cross-Linking Immunoprecipitation(Chi et al., 2009) technique in the retina rather than rely on computational prediction algorithms (Chi et al., 2009, Moore et al., 2014). HITS-CLIP, here using the Argonaute-2 protein (AGO2) as bait, enabled high confidence validation of miRNA targets which has historically been problematic when based on computational *in silico* predictions (Jensen and Darnell, 2008, Licatalosi et al., 2008, Ule et al., 2005, Ule et al., 2003, Yeo et al., 2009). AGO2 as a core component of the RISC complex has been implicated in the biogenesis and maturation of miRNAs, in addition to its function of directly binding mature miRNAs for mRNA target repression (Petri et al., 2014). Specifically, CLIP of AGO2 enables the identification of only those miRNAs, that are currently complexed within RISC and thus biologically functional, and their resultant targetome (Jensen and Darnell, 2008, Licatalosi et al., 2008, Ule et al., 2005, Ule et al., 2003, Yeo et al., 2009)

Our previously published transcriptome-wide miRNA and mRNA comparison using standard small RNA and RNA high-throughput sequencing (HTS) in a focal, mouse photo-oxidative damage (PD) model of retinal degeneration defined the retinal miRnome (Natoli et al., 2016). This confirmed the involvement of pathways that mimic the key pathological features of AMD such as inflammation dysregulation and complement activation. In the study presented here, using AGO2 HITS-CLIP, we revealed that only a small miRNA subset made up the majority of the retinal miRnome with miR-124-3p being highly represented in AGO2-bound miRNA. While the retinal AGO2-bound miRnome did not differ between healthy and damaged retinas, a shift was observed in the mRNA targetome following retinal damage that was associated with a dynamic change in seed region binding specifically of miR-124-3p.

Finally, we showed that AGO2 accumulated in the inner retinal layers, such as the glial cells, upon retinal damage, which potentially signifies a major shift in miRNA activity in these cells. Together these results provide mechanistic insight into the alteration of miRNA-based transcriptome regulation, offering an understanding of how this contribution to the degenerating retina could help pave the way for effective miRNA-based therapeutics for the treatment of retinal degenerations including AMD.

## Results

### Global miRNA and mRNA transcriptomes reveal key miRNA and inflammatory pathways involved in retinal degeneration induced by PD

High throughput sequencing (HTS) of miRNA and mRNA was initially performed globally (no Ago immunoprecipitation) to decipher overall changes in the retina between dim-reared (control, DR) and photo-oxidative damage (PD) mouse retinas. 3-9 million and 47-71 million cleaned reads were generated from miRNA and mRNA samples, respectively. Significant homogeneity was seen between the samples within each group for both mRNA (Figure 1A, blue) and miRNA (Figure 1A, green), indicating high reproducibility of data. mRNA analysis of the global retinal HTS identified 3018 mRNA transcripts to be significantly changed between DR and PD retinas based on a p-value of less than 0.05 (Figure 1B). Gene ontology (GO) and pathway analyses of these differentially expressed mRNA revealed a number of significantly enriched terms/pathways involved in inflammation including complement activation, positive regulation of interleukin-1 production, macrophage activation, innate immune response and response to cytokine (Figure 1C). 287 differentially expressed miRNA were identified when comparing the DR and PD groups (FDR<0.05). The top 80 differentially expressed miRNA showed high intra-group similarities, but significantly different inter-group differences between DR and PD groups (Figure 1D). Among the top 10 most significantly changed miRNA were those known to be involved in maintaining retinal neuronal homeostasis (Karali and Banfi, 2018) including miR-9-5p (down-regulated), miR-211-5p (down-regulated) and miR-191-5p (up-regulated) (Figure 1E). Interestingly, the top 20 global expressed miRNA represented 85% of the total retinal miRnome in both the DR (Figure 1F) and PD (Figure 1G) retinas, with let-7 family members, photoreceptor-enriched miR-183-5p and miR-182-5p, and neuron-enriched miR-124-3p the most highly expressed miRNAs in both groups. Together, these results demonstrate the prominence of the innate immune system in the progression of retinal degeneration in our PD model and that only a subset of miRNA represents an overwhelming majority of the total retinal miRnome with transcripts from our analysis.

**Figure 1.**
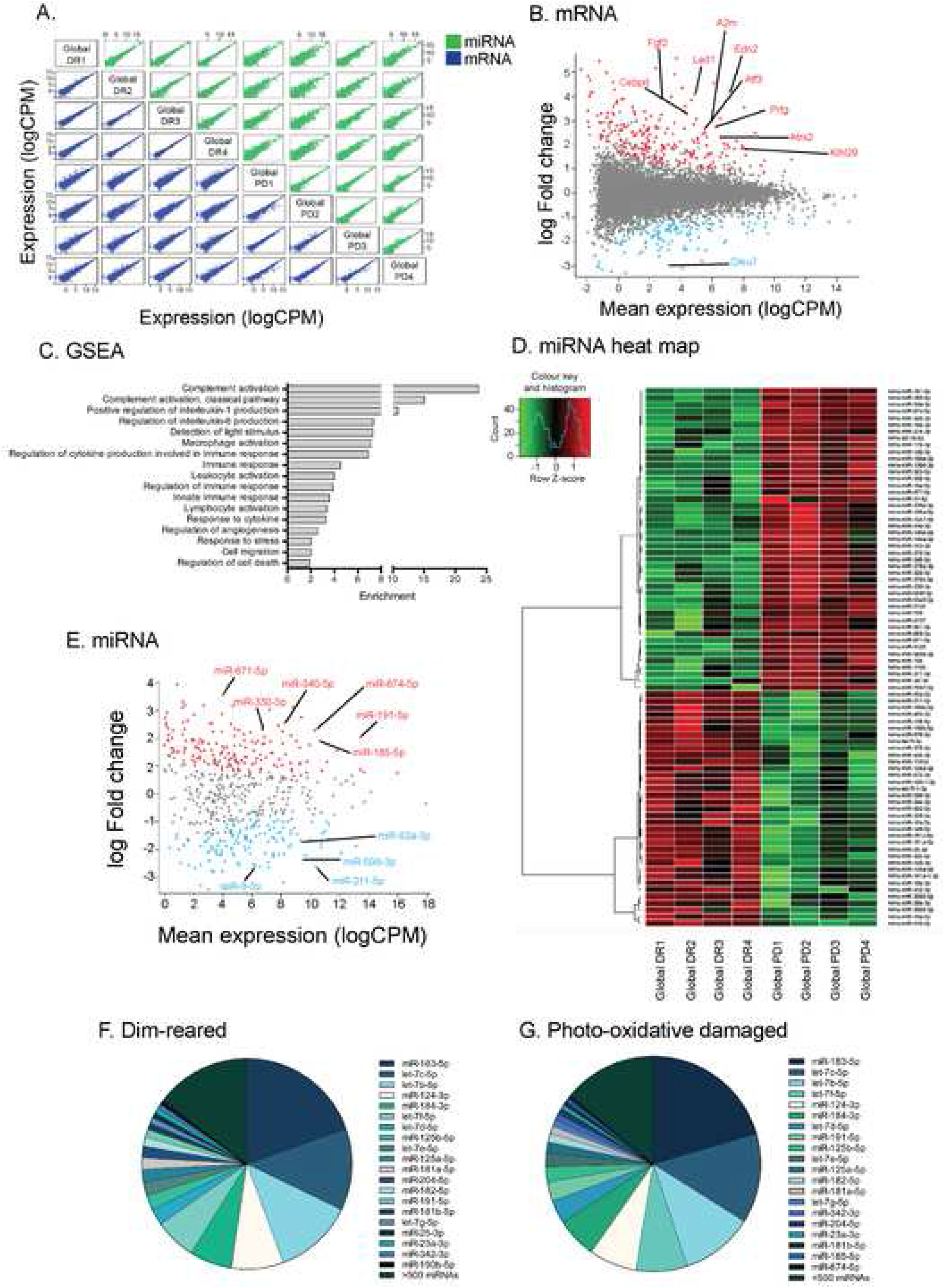
Global mRNA and miRNA identification in the retina by high throughput sequencing. (A) Visualisation of intra- and inter-group variation with a correlation plot comparing log CPM of the global mRNAs (blue) and miRNA (green) identified between DR and PD samples. Each dot in the figures represents a gene expression (log2(count per million)) in samples labelled in diagonal panels. Homogeneity was seen between intra-group samples with variation in inter-group samples indicating differential expression. (B) Mean-difference plots of global retinal mRNA with significantly up- and downregulated transcripts shows in red and blue, respectively (P<0.05). The top 10 most significantly changed transcripts are individually labelled (FDR<0.05). (C) GSEA analysis of the differentially expressed mRNA transcripts revealed a number of significantly enriched terms involved in inflammation and the immune system. (D) Hierarchical clustering of the top 80 differentially expressed miRNA showed significant homogeneity between intra-group samples. (E) Mean-difference plots of global retinal mRNA with significantly up- and downregulated transcripts shows in red and blue, respectively (P<0.05, FDR<0.05). (F-G) Pie chart showing the proportions of the globally expressed miRNA in the miRnome for dim-reared (F) and photo-oxidative damage (G) retinas.

### AGO2 HITS-CLIP reveals retinal miRnome and targetome

Whilst the global transcriptome analysis provides a snapshot of the molecular and physiological processes that occur in response to PD, AGO2 immunoprecipitation was performed on DR and PD retinal samples to decipher the functional miRnome and its respective targets (Figure 2A). Initial analyses confirmed the specific immunoprecipitation of AGO2-bound radiolabelled retinal miRNAs and mRNAs with two distinct populations visible through gel electrophoresis: 1) binary AGO2:miRNA complex resolves at 110 kDa; and 2) a ternary AGO2:miRNA:mRNA complex “smear” at 130 kDa (Chi et al., 2009, Moore et al., 2014) (Figure 2B). To improve RNA end yield, miRNA/RNA extraction kits were utilised for AGO2 HITS-CLIP (Figure 2C). Total AGO2-bound RNA was extracted and separated into mRNA (Figure 2D) and small RNA (including miRNA, Figure 2E) fractions through a filtration process (miRvana, Thermo Fisher Scientific) before sequencing.

**Figure 2.**
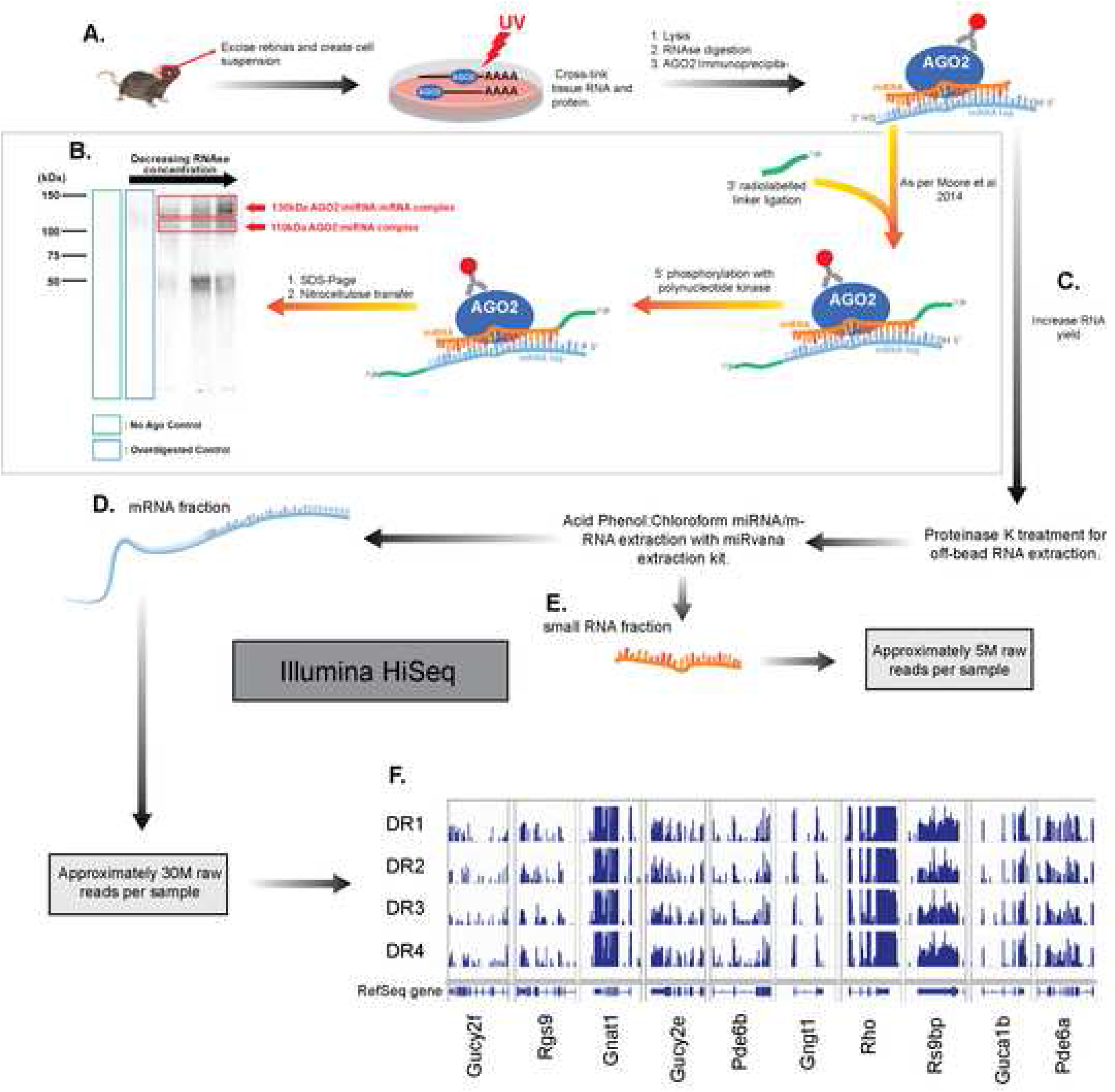
Workflow of AGO2 HITS-CLIP in mouse retina. (A) Retinas were excised and diced into a fine suspension before following a series of steps for immunoprecipitation (IP) as described in detail in the methods. (B) AGO2 IP was performed as described previously (Moore et al. 2014). Phosphorimage of gel electrophoresis of end-labelled RNAs purified by AGO2 IP showed 110kDa and 130kDa bands representing the AGO2:miRNA and AGO2:miRNA:mRNA complexes, respectively. A no AGO2 antibody (green box) and overdigested control (excess RNase, blue box), were also included. (C) RNA and miRNA extractions following HITS-CLIP were performed to separate both fractions before Illumina HiSeq, which yielded approximately 30 million reads for the mRNA fraction (D) and 5 million reads for the miRNA fraction (E). (F) Read coverage (blue bars) of the four DR biological replicates for genes in the rhodopsin pathway displayed high reproducibility. The mm10 gene annotation according to RefSeq is shown at the bottom as horizontal blue bars, with gaps indicating intronic regions.

To determine the size and breadth of the predicted target networks of the DR retina AGO2-bound miRNA, miRNet network analysis was performed for the top 20 miRNA (present in AGO2:miRNA complexes). This analysis revealed mRNA target networks for 14 of the miRNA (Figure 3A) with the other six not well annotated in the system. Among them, miR-124-3p and miR-181-5p returned the majority of gene target interactions with 223 and 233 interactions each, respectively (Figure 3A). We identified 553 AGO2-bound miRNAs in DR retinas. Interestingly, miR-124-3p was overwhelmingly expressed in DR retinas, representing approximately 75% of the AGO2-bound miRnome (Figure 3B). Photoreceptor specific miR-183-5p (part of a cluster miR182/96/183) (Karali et al., 2010, Karali et al., 2007, Krol et al., 2010, Xiang et al., 2017, Xu et al., 2007) was the second most abundant miRNA followed by miR-124-5p. The remaining top 10 included miR-125b-5p, miR-125a-5p, miR-181a-5p, miR-191-5p and three members of the miR-29 family: miR-29a-3p; miR-29b-3p; and miR-29c-3p. The two other arms of the photoreceptor cluster, miR-182-5p and miR-96-5p, were also among the top 20 AGO2-bound miRNA in DR retinas. miR-301a-3p, miR-130a-3p, miR-204-5p, miR-99b-5p, miR-29b-1-5p, miR-342-3p, with two members of the let-7 family (let-7d-3p and let-7f-5p) completed the top 20 (Figure 3B). A total of 9397 mRNAs were identified as bound to AGO2 in DR retinas.

**Figure 3.**
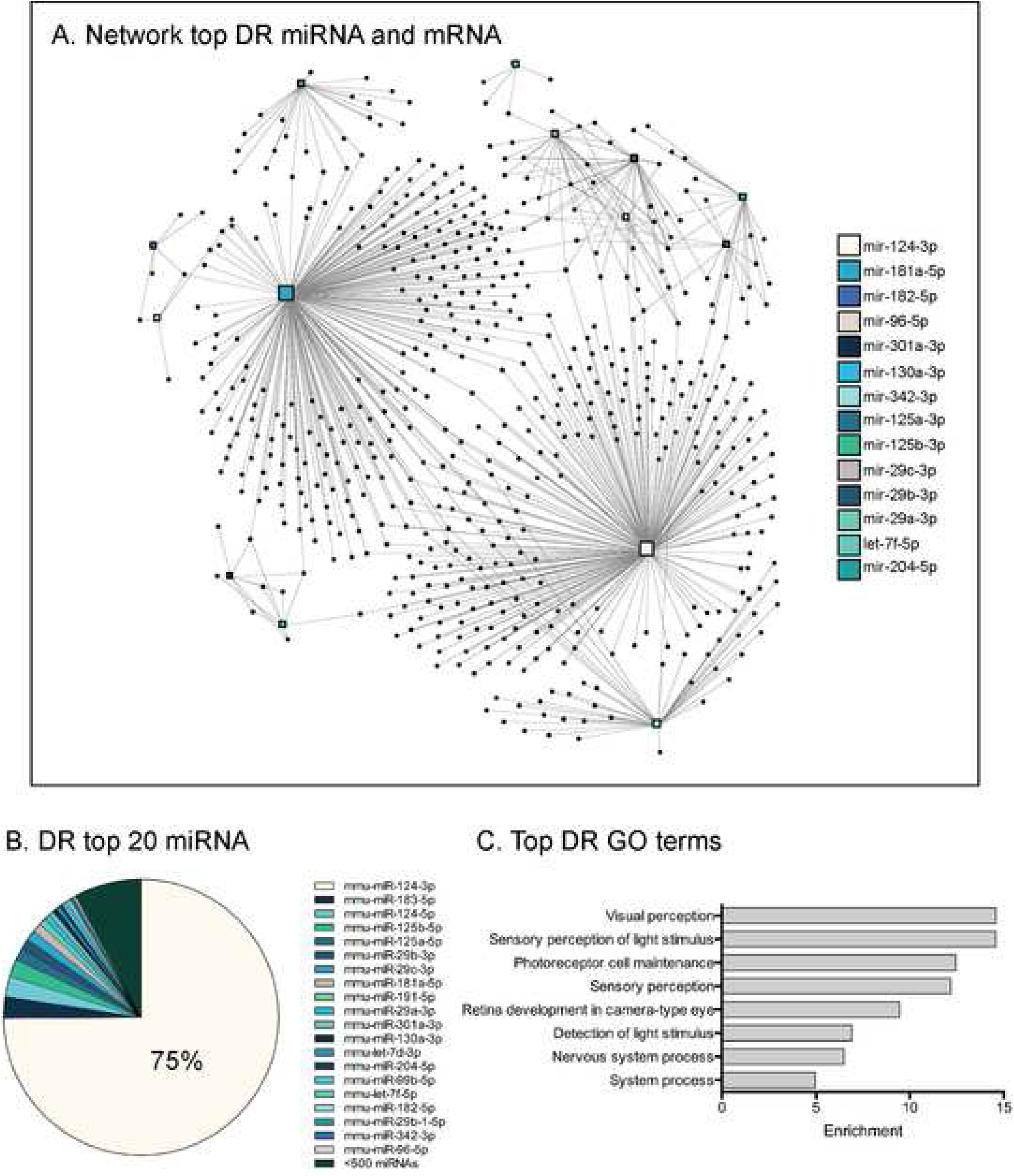
Interaction network analysis of the top 20 DR retinal AGO2-bound miRNA. (A) Illustration the network analysis results for 14 miRNA against all AGO2-bound mRNA transcripts, with miR-124-3p and miR-181a-5p having the largest amounts of connections. The remaining six miRNAs were not annotated in the system and did not yield a network. Each dot represents an interacting mRNA connected by lines to the respective miRNA. The size of the centre square indicates number of mRNA interactions for each miRNA. (B) Pie chart of the AGO2 miRnome of DR retinas, miR-124-3p constitutes approximately 75% of the AGO2 miRnome. (C) Gene ontology analysis was performed based using the most abundant AGO2-bound mRNA transcripts in DR retinas (P<0.05 considered significant).

GO analysis of the top DR AGO2-bound mRNA transcripts revealed significant enrichment in processes involved in vision such as visual perception, sensory perception of light stimulus, photoreceptor cell maintenance and detection of light stimulus (P<0.05, Figure 3C).

Here we have demonstrated effective isolation of both AGO2:miRNA binary and AGO2:miRNA:mRNA ternary complexes. These complexes contain a small subset of miRNA that constitute the active retinal miRnome during homeostasis, with miR-124-3p the most abundant in the population. Further, the functional miRnome is involved in standard visual processes of the retina, indicating a homeostatic role for vision and retinal maintenance.

### Retinal AGO2-bound miRNA shifts targetome following PD

Similarly, miRNet network analysis of the top 20 AGO2:miRNA in PD samples also showed networks for 14 of the AGO2-bound miRNA (Figure 4A). Again, as for the DR retinas, miR-124-3p and miR-181-5p displayed the highest number of interactions (Figure 4A). miR-124-3p was also the most abundant AGO2-bound miRNA in PD retinas comprising 81% of the miRnome (Figure 4B). Similar to the DR retinas, miR-124-5p and miR-183-5p were second and third highest expressed, respectively. The remaining top 10 AGO2-bound miRNA were similar to that found in DR retinas with the miR-125 family, two members of the miR-29 family, miR-181a-5p, mir-191-5p and miR-130a-3p identified. The highly enriched GO terms for the top PD AGO2-bound mRNA were similar to those in the DR samples, such as visual perception and sensory perception of light stimulus (Figure 4C).

**Figure 4.**
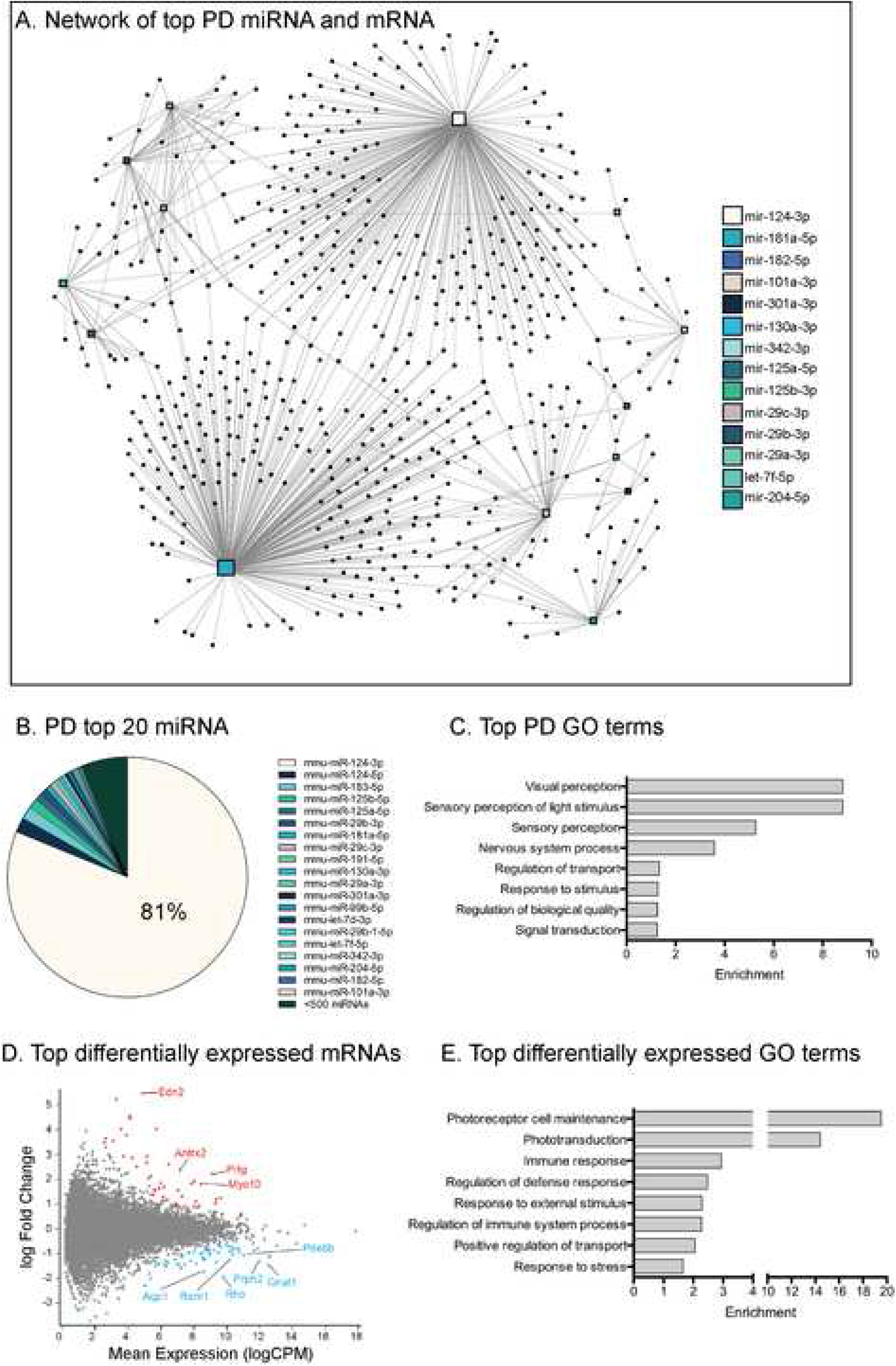
Interaction network analysis of the top 20 PD retinal AGO2-bound miRNA. (A) Illustration of the network analysis results for 14 miRNA against AGO2-bound mRNA transcripts, with miR-124-3p and miR-181a-5p having the largest amounts of connections. The remaining six miRNAs were not annotated in the database and yielded no result. (B) Pie chart of the AGO2 miRnome of PD retinas, with miR-124-3p constituting approximately 81% of the miRnome. (C) Gene ontology analysis was performed using the most abundant AGO2-bound mRNA transcript in DR retinas. (D) Mean difference plot of the top 10 most differentially AGO2-bound mRNA between DR and PD (FDR<0.05). Significantly over- and under-represented mRNA are shown in red and blue, respectively (P<0.05; FDR<0.05). The top 10 most significantly different mRNA are labelled individually (P<0.05). (E) Gene ontology analysis was performed based on the most significant differentially expressed Ago mRNA transcripts between DR and PD retinas. This revealed a number of enriched terms involved in inflammation and the immune response (P<0.05 considered significant).

Although AGO2-bound mRNA enriched GO terms between DR and PD samples were similar, we asked whether there were any differences. We found *Gnat3* and *Rho* to be among the top 10 most differentially bound mRNAs, two mRNAs involved in the process of photoreceptor phototransduction, the process that converts entering light into an electrical signal (Figure 4D). Interestingly, when conducting GO analysis using the significantly differentially AGO2-bound mRNA transcripts, the enriched terms related to inflammation such as immune response, regulation of defence response, regulation of immune system process, and response to stress (Figure 4E). Further gene set enrichment analysis (GSEA) against the C2 database resulted in specific pathways such as complement, NF-KB and EGF signalling pathways differentially expressed between AGO2-bound PD and DR retinas (Supplementary Table 1). These results indicate that whilst maintenance of the visual system remains a priority for the retinal miRnome, there is a nuanced change in mRNA targeting by functional AGO2-mediated miRNA following photo-oxidative damage.

### miR-124-3p undergoes shift in seed binding regions following PD

Given the changes of targeted mRNA following PD, we next analysed the binding interactions of the AGO2-bound miRNA with their targetome. We performed peak calling of the 3531 unique AGO2-bound mRNA transcripts identified with HITS-CLIP in DR and PD retinas. The most enriched sequence motif within the 3’UTR in DR retinas was the 6-mer (reverse complement of miRNA positions 2-7) and 7A1 (reverse complement of miRNA positions 2-7 with an adenine base) seed sequences for miR-124-3p (Figure 5A). In PD retinas we found that in addition to the 6-mer and 7A1 motifs, the 7m8 (reverse complement of miRNA positions 2-8) and 8-mer motifs (reverse complement of miRNA positions 2-8 with an adenine base) were additionally present (Figure 5B). Amongst the mRNA that were differentially AGO2-bound in PD retinas, the transcripts predicted to contain the 7m8 (Figure 5C) and 8-mer (Figure 5D) seed sequences of miR-124-3p were graphed. A number of these transcripts have well-established links to retinal degeneration including chemokine C-C motif ligand 2 (*Ccl2*) and rho-associated protein kinase 2 (*Rock2*) (Figure 5D, P<0.05). The full list of AGO2-bound mRNA targets of miR-124-3p are presented in Supplementary Table 2.

**Figure 5.**
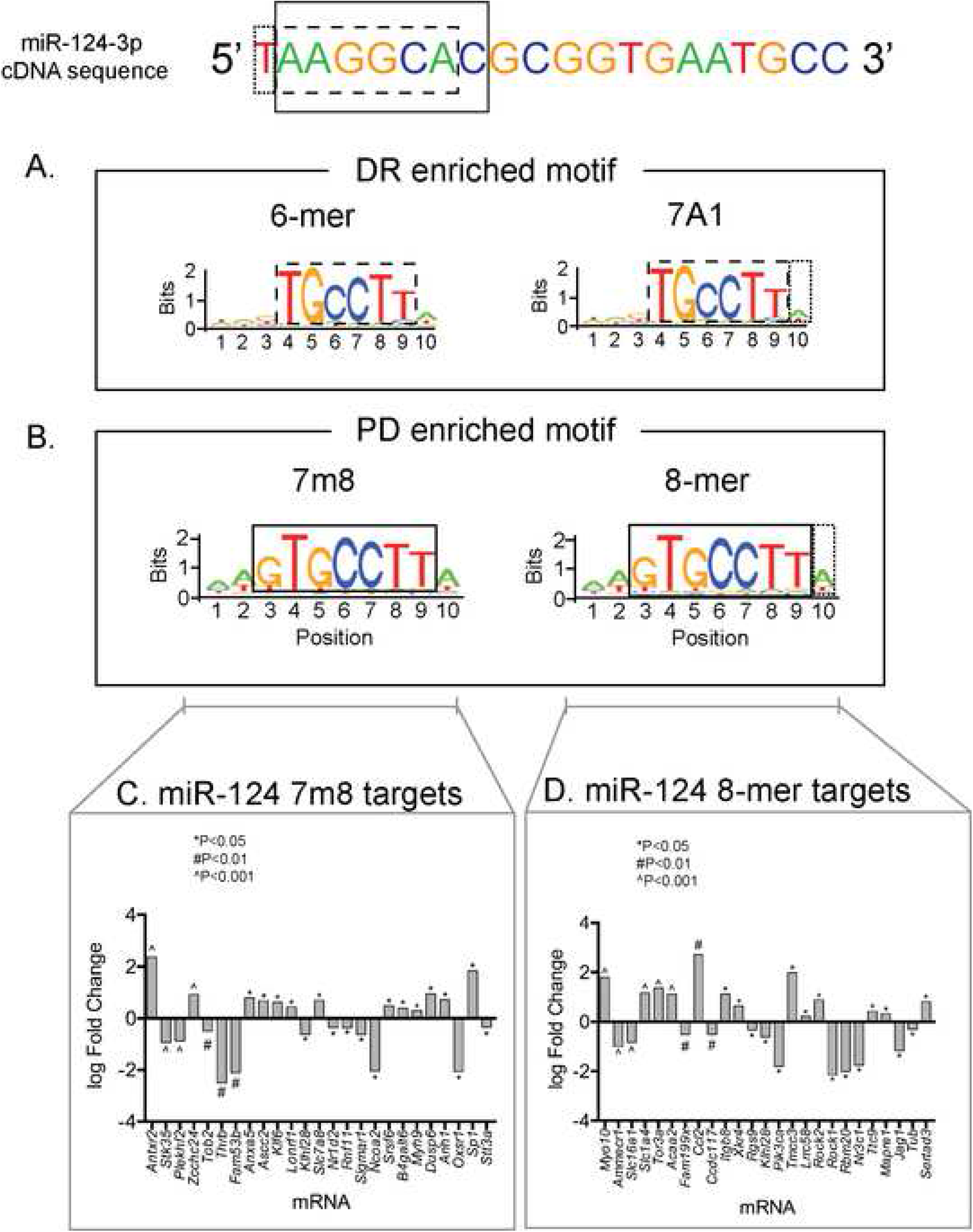
Enriched sequence motifs of AGO2-bound mRNA. The RNA sequence of miR-124-3p is displayed at the top of the figure. (A) Most enriched binding motif of DR retinas were the 6-mer (dashed line) and 7A1 motifs (reverse complement of positions 2-7 of the miRNA (dashed line), and with an additional adenine base (dotted line), respectively). (B) Most enriched binding motif of PD retinas, in addition to the 6-mer and 7A1, were the 7m8 (solid line) and 8-mer motifs (reverse complement of positions 2-8, and with an additional adenine base (dotted line), respectively). (C) 25 AGO2-bound mRNA had complementarity with the 7m8 seed sequence. Similarly, 25 AGO2-bound mRNAs had complementarity to the 8-mer seed sequence. (^P<0.001, #P<0.01, *P<0.05).

Since it has been reported that miR-124-3p may shift from a neuronal to a glial expression profile following PD (Chu-Tan et al., 2018), we performed a literature search for known cellular locations of the AGO2-bound mRNA targets of miR-124-3p described above (Supplementary Table 2). Approximately 29% of AGO2-bound miR-124-3p targets in PD have been described to localise to the INL of the retina with a further 23% localised to the GCL (Figure 6A). Amongst those genes reported to be expressed in the INL, 80% were located specifically within the Müller glial cells (Arnold et al., 2012, Mohammad et al., 2018, Rutar et al., 2011, Sarthy et al., 2005, Soundararajan et al., 2014, Ueki et al., 2015). The expression level of these 8 Müller glial genes increased in PD retinas, as determined here by AGO2 HITS-CLIP of DR and PD retinas, with the exception of *Rock1* (P<0.05, Figure 6B).

**Figure 6.**
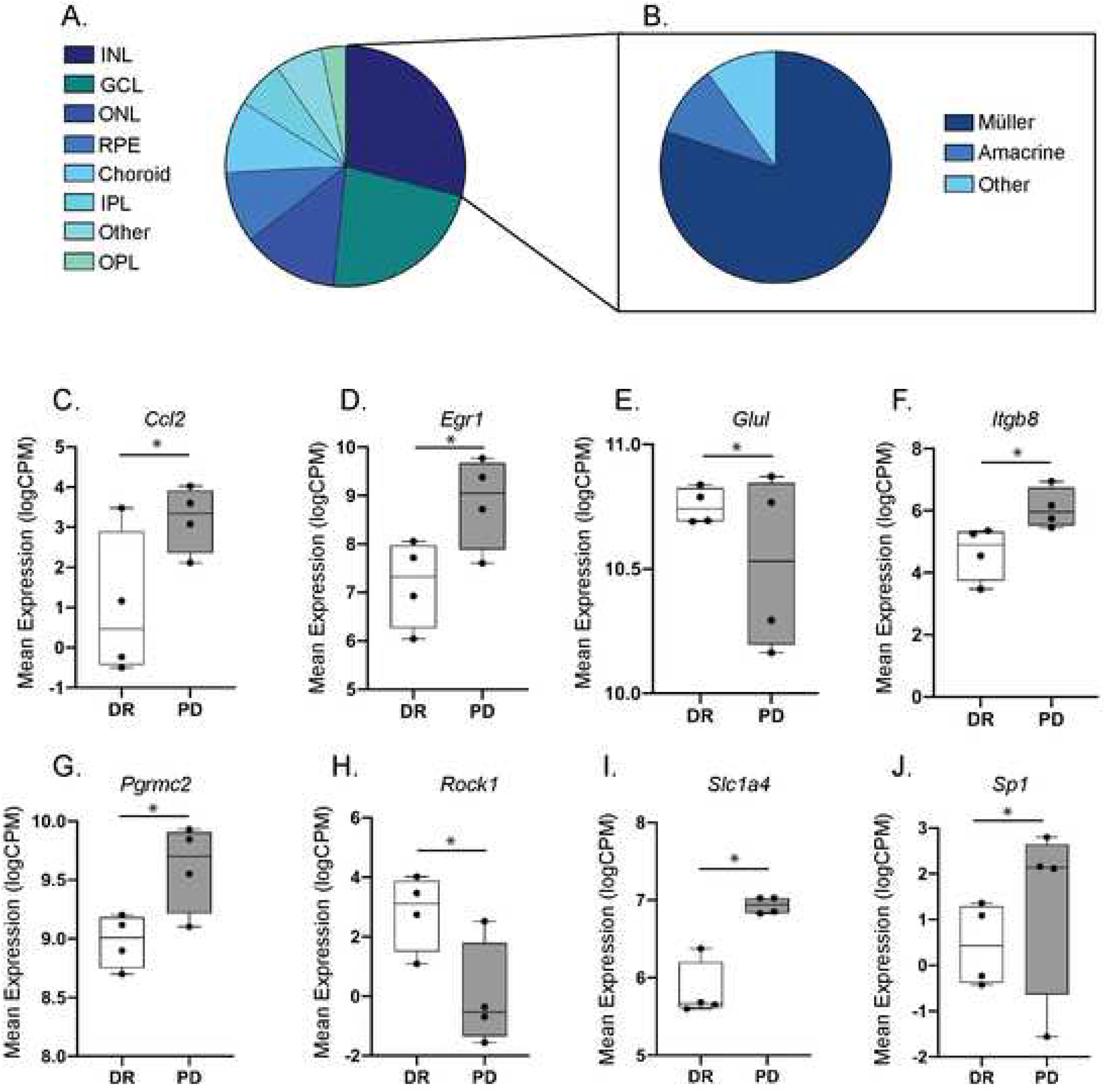
Characteristics of AGO2-bound mRNA targeted by miR-124-3p. (A) Pie chart of the published cellular localisation of AGO2-bound miR-124-3p targeted mRNA transcripts shown in Figure 5C/Supp Table 2. Most were located to the INL and GCL. 80% of INL localised transcripts were located specifically to Müller glial cells. (B) The expression level of the Müller glial cell transcripts showed a general trend of higher expression in PD retinas compared to DR (*P<0.05).

### Retinal AGO2 increases in the inner neuronal retinal layers following PD

In order to provide a measure of spatial information to associate with the HITS-CLIP data and to tie in the miRNA seed sequence data, the relative retinal mRNA expression levels of *Ago2* and its protein localisation in the retina was investigated using both DR and PD retinal cryosections and tissue lysates. *In situ* hybridisation of *Ago2* using DIG-labelled riboprobes, revealed an increase in *Ago2* labelling specifically in the GCL and INL of the retina following PD when compared to DR retinas, in which minimal labelling is seen (Figure 7A). Immunohistochemistry of AGO2 showed a similar labelling pattern, with high AGO2 expression seen in the GCL and INL as well as in the OLM and outer segments of the photoreceptors in PD retinas when compared to DR retinas (Figure 7B). Furthermore, using qRT-PCR, *Ago2* expression levels were found to be significantly increased in PD retinas compared to DR controls (Figure 7C, P<0.05). Western blot quantification of AGO2 protein abundance demonstrated a significant increase in PD retinas when compared to DR after normalising to the reference protein GAPDH (Figure 7D and 7E, P<0.05). These results strongly indicate that AGO2 expression increases and accumulates in the inner retinal layers, potentially localised to Müller glia, suggesting a concerted effort to direct miRNA activity to cells in these regions following PD.

**Figure 7.**
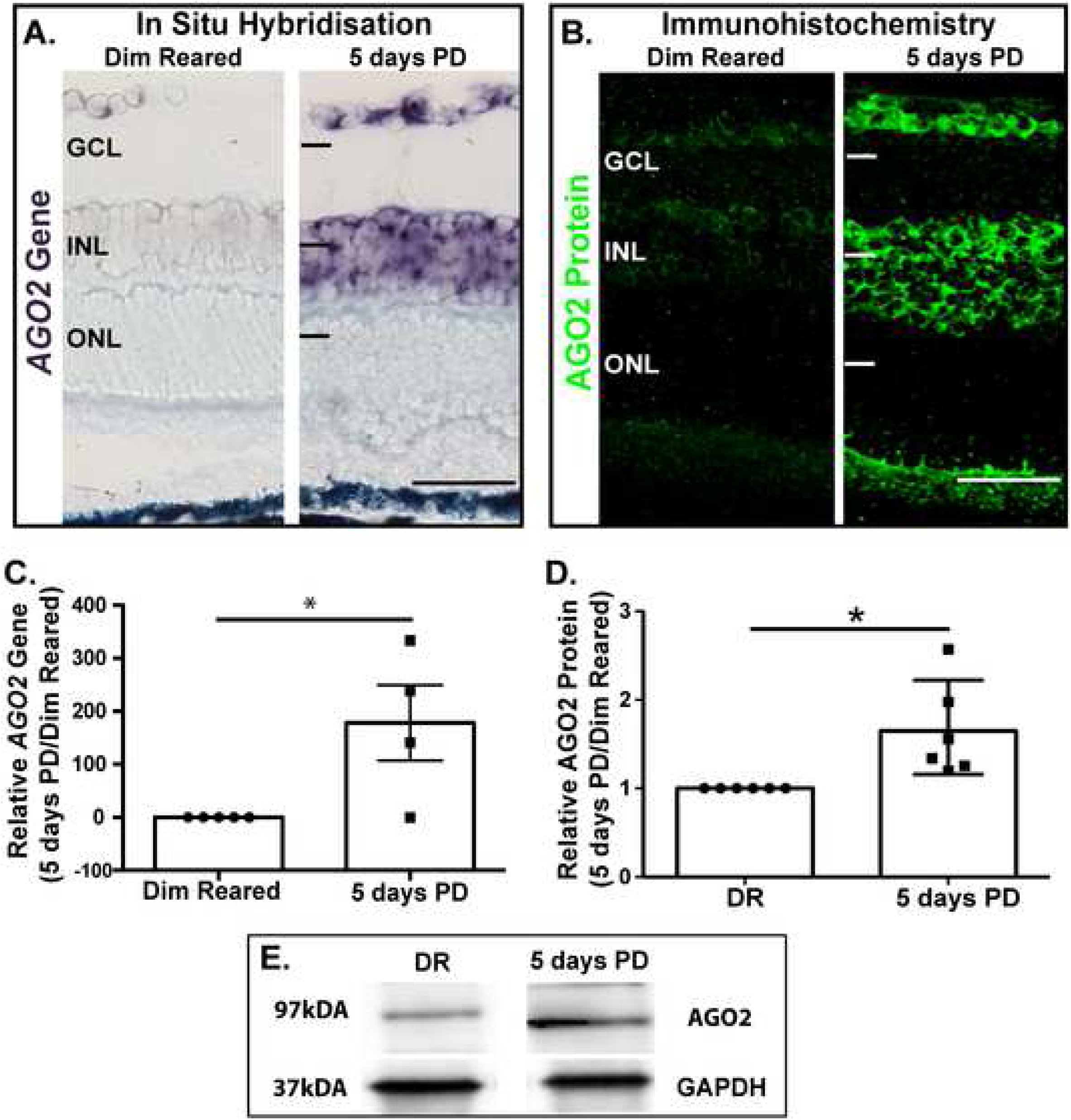
AGO2 expression profiles in retinal histology. (A) *In situ* hybridisation of *Ago2*. *AGO2* mRNA expression increased dramatically following 5 days of PD when compared with DR retinas, primarily in the INL and GCL. (B) Immunohistochemistry labelling AGO2 protein. AGO2 protein displayed the same trend as its mRNA but with additional labelling in the outer segments of the photoreceptors or end feed of the Müller glial cells. (C) qRT-PCR revealed a significant increase in *Ago2* expression in PD retinas when compared to DR. (D) Optical densitometry of the immunohistochemical staining of AGO2 resulted in a higher relative expression level in PD retinas compared to DR. (E) Western blot of AGO2 protein also indicated an increase in expression level in PD retinas (*P<0.05 Student’s *t*-test).

Taken together, these results suggest a change in miRNA target activity with a possibility of increased movement to glial cells following PD. These changes may reflect a key aspect of miRNA behaviour under duress that leads them to retinal maintenance cells, such as glia, to regulate differentially expressed disease-specific pathways in the retina.

## Discussion

miRNA are the most commonly studied non-coding RNA in the central nervous system (CNS) (Cao et al., 2016), and the retina (Karali and Banfi, 2018). Despite this, the specific miRNAs required to maintain retinal homeostasis or response to retinal degeneration have not been identified. Studies identifying the retinal miRnome under homeostatic conditions have utilised small RNA sequencing technology to investigate the total present miRNA with high resolution and sensitivity (Karali et al., 2016). Our study extends this data by utilizing an AGO2 HITS-CLIP method thus allowing for the identification of not only the miRNA, but also the functionally active mRNA that is bound in a AGO2:miRNA complex *in vivo.* We demonstrated that both global miRNA and mRNA expression are regulated in response to PD, and that the inflammatory and cell death gene pathways are highly enriched. Furthermore, our data characterise the functional retina miRnome during normal conditions (DR) and retinal degeneration (PD). Interestingly we observed that the AGO2-bound miRNAs did not change in response to PD but that the AGO2-bound mRNA targets differed. Importantly the AGO2-bound miRNAs and mRNAs have been linked to key pathways already identified as important in human AMD pathogenesis, such as those that regulate the immune response. Finally, our data demonstrated that following PD the expression of the AGO2 gene (mRNA) and protein was upregulated in the INL and GCL of the retina. We suggest that this upregulation of AGO2 indicates a potential glial stress response to retinal degeneration, and thus could be mediated by a change in AGO2-miRNA activity by directing miRNA to areas of need. This hypothesis was further corroborated with seed enrichment analysis demonstrating a change in the seed region binding activity of the highly expressed miR-124-3p following PD, with a number of PD-specific AGO2-bound targets known to be localised within retinal glial cells. These data overall demonstrate a dynamic role of the retinal miRnome with its targetome, which is influenced by the state of the tissue environment providing important insight into miRNA functionality during neurodegeneration.

We used standard HTS of the retina to provide a snapshot of the miRNA and biological pathways at play in the healthy retina and in retinas subject to PD-induced retinal degeneration. Many reports have supported the notion that the vast majority of miRNAs in a cell are comprised of only a select few highly abundant miRNAs (Couzin, 2008, Meola et al., 2009, Sanuki et al., 2011, Wienholds et al., 2005). This suggests that these particular miRNAs may hold key roles in gene regulation of any given system. This holds true in the retina, with results from this study revealing that approximately 85% of the entire retinal miRnome consists of only 20 different miRNA in both DR and PD retinas. Included in this group are members of the photoreceptor cluster miR-183/96/182, which has long been documented to be involved in the development and homeostasis of photoreceptor cells. Further, miR-124-3p was also highly expressed as a neuronally-enriched miRNA that plays a large role in the maintenance of neuron homeostasis (Conaco et al., 2006, Lagos-Quintana et al., 2002, Lim et al., 2005, Makeyev et al., 2007).

Expectedly, AGO2-loaded miRNA found in both DR and PD groups, comprised miRNAs that are known to play a role in retinal development and homeostasis such as the photoreceptor cluster miR-183/96/182 (Bellon et al., 2017, Busskamp et al., 2014, Fan et al., 2017, Karali et al., 2007, Krol et al., 2010, Lumayag et al., 2013, Xiang et al., 2017, Xu et al., 2007); miR-204-5p (Barbato et al., 2017, Conte et al., 2010, Conte et al., 2015, Karali et al., 2016), let-7a-5p (La Torre et al., 2013) and miR-124-3p (Chu-Tan et al., 2018, Karali et al., 2007, Sanuki et al., 2011), showing consistent overlap with previous studies. However, despite significant changes in differential global miRNA expression in the damaged retina, there were no significant changes in AGO2-loaded miRNA between DR and PD retinas suggesting that the retina appears to be utilising the existing miRNA population in both physiological states. What is novel, however, is that despite similarities between the DR and PD AGO2-bound retinal miRnomes, AGO2-bound mRNA (the targetome) displayed significant differences between the control and damaged states. With AGO2-loaded miR-124-3p and miR-181a-5p (yet another highly-enriched retinal miRNA (Carrella et al., 2015a, Carrella et al., 2015b)) displaying extensive mRNA target networks in the population, this finding is again suggestive that only a set of highly enriched miRNAs are responsible for the regulation of a dynamic group of mRNA targets (Helwak et al., 2013, Jalali et al., 2013, Li et al., 2014, Paraskevopoulou et al., 2013, Schug et al., 2013, Ziu et al., 2014).

Based on the results from GO term and pathway analyses of the global mRNA sequencing, it was revealed that photoreceptor maintenance and visual cycle were highly enriched in both DR and PD states. This was also true when we looked at the AGO2-bound mRNA in both DR and PD in terms of mRNA abundance indicating, perhaps unsurprisingly, that under both states maintenance of the visual system still remains a regulatory priority in the retina. However, the global HTS analysis also revealed that many of the enriched PD pathways were involved in retinal inflammation (Ambati et al., 2013). The GO terms and pathways for the differentially expressed AGO2-bound mRNA demonstrated involvement in pathways of the immune response, regulation of defence, regulation of immune system processes, and response to stress. These pathways have been strongly implicated in the progression of retinal degeneration which others and we have reported (Edwards et al., 2005, Hageman et al., 2005, Haines et al., 2005, Klein et al., 2005, Natoli et al., 2016).

Highly represented is the complement pathway, which has been previously identified as a major instigator of the inflammatory dysregulation that occurs in AMD pathogenesis (Edwards et al., 2005, Hageman et al., 2005, Haines et al., 2005, Klein et al., 2005). The immune response, particularly the innate immune response, is well-established to influence the progression of retinal degenerations (Ambati et al., 2013). Pathways involved in the innate system such as the interleukin and cytokine release were also significantly enriched in the differentially expressed dataset. This is suggestive of a glial response to stress as glial cells such as Müller glia, play a large role in both the maintenance and contribution to retinal inflammation. With a firm understanding that these processes and pathways are highly represented in our model of retinal degeneration, determining the miRNA binding sites within these genes along with their regulatory miRNA using AGO2 HITS-CLIP, may lead to advances in potential therapeutic options targeting key inflammatory components.

Determination of binding sites is made possible with seed motif enrichment analysis which, when applied to a dataset, can elucidate enriched binding motifs of the AGO2-loaded miRNA. We demonstrated that the most enriched seed region motif in PD was the canonical 7m8 and 8-mer seed regions of miR-124-3p, but interestingly DR retinas had a different enriched region (6-mer and 7A1) within the seed sequence, which suggests a different manner of binding under stress. This change in seed region may reveal that miRNA regulation actually acts under the influence of changes in stimuli in their environment (Nigita et al., 2016).

We concentrated our data analysis on the binding targets of miR-124-3p due to the observed change in seed region, its abundance in both the DR and PD results, and its known role in retinal degenerations (Chu-Tan et al., 2018). Particularly interesting was the expression of *Ccl2* as it was among the AGO2-bound miR-124-3p targets significantly increased in PD retinas with a seed region (8-mer) not enriched for in healthy DR controls. This biological interaction between miR-124 and *Ccl2* has been demonstrated to increase in AMD patient retinas (Newman et al., 2012), and is one of the integral signalling molecules required to recruit monocytes to the site of retinal damage which exacerbates degeneration (Rutar et al., 2015, Rutar et al., 2012, Rutar and Provis, 2016). Further regulation of CCL2 activity has been shown to slow retinal degeneration, with increased chemokine signalling postulated as a key instigator to the dysregulated immune response seen in AMD (Newman et al., 2012, Rutar et al., 2015, Rutar et al., 2012, Rutar and Provis, 2016, Guo et al., 2012, Sennlaub et al., 2013). Our previous studies investigating miR-124 and *Ccl2* interactions indicated a potential cellular translocation of miR-124 from neuronal to glial, largely Müller cells (Chu-Tan et al., 2018). Approximately 25-30% of the AGO2-bound mRNA targets of miR-124-3p identified in this study have previously been identified to be located to retinal Müller glial cells (Arnold et al., 2012, Mohammad et al., 2018, Rutar et al., 2011, Sarthy et al., 2005, Soundararajan et al., 2014, Ueki et al., 2015). In addition to this, the expression of these targets seem to be increased after PD suggesting a potential movement of miRNA activity to Müller cells upon stressed conditions, consistent with the finding of miR-124 and *Ccl2* previously published by our group (Chu-Tan et al., 2018).

Taken together with results from this work, we postulate that miRNA activity in the damaged retina is differentially regulated compared to the homeostatic DR state, however, does not directly change the relative loading levels of the miRNA themselves into AGO2. It is likely that this demonstrates a system in which cells utilise the existing population of miRNA to regulate different targets, rather than transcribe novel miRNAs, an idea that has been postulated before (Nigita et al., 2016) but yet to be shown in the retina. So, whilst the primary miRNA remain consistent in their overall activity, a shift in their interactions represents a change in the regulatory priorities of the entire system. This dynamic change may represent an early response to disease progression that may eventually lead to increased loss in miRNA activity, a hypothesis that requires further testing using an extended damage paradigm.

An alternative explanation to these differences observed could be that the canonical pathway of miRNA biogenesis might not be the only pathway contributing both to retinal homeostasis and retinal degenerations. The data generated from these experiments will be useful in determining alternative, non-canonical binding sites for the miRNAs of interest, which is afforded through use of the HITS-CLIP methodology. Work in the field is beginning to shift to focus on miRNA binding events that occur outside of the 3’UTR (Boudreau et al., 2014) which would help explain the difference of gene expressions seen in the HTS global sequencing experiments compared to the HITS-CLIP analysis. Further analysis of the enriched peaks in our data may reveal binding sites in the coding regions, 5’UTR, intronic regions, and wobble or bulge nucleotides, which will be indicative of non-canonical targets that can be explored further (Didiano and Hobert, 2006, Tay et al., 2008, Vella et al., 2004). Future work should address the functional relevance of these binding sites and the targets associated with them in order to fully uncover the functional miRnome in retinal degenerations.

Finally, results from this work indicate a possible shift in miRNA activity from largely neuronal to glial following damage (Figure 8). We investigated the expression profile and location of AGO2 in order to indirectly determine where this change in AGO2-bound miRNA activity may occur in the retina. In addition to being the binding “vehicle” for miRNA function, removal of mature miRNAs through *Dicer* or *Dgcr8* knockouts, consequently reduces AGO2 levels (Gibbings et al., 2012), further demonstrating the closely interconnected relationship between mature miRNA and AGO2. We showed that the retinal AGO2 protein and *Ago2* gene increased after PD and localised to the INL and GCL, two of the retinal layers spanned by Müller glial processes.

**Figure 8.**
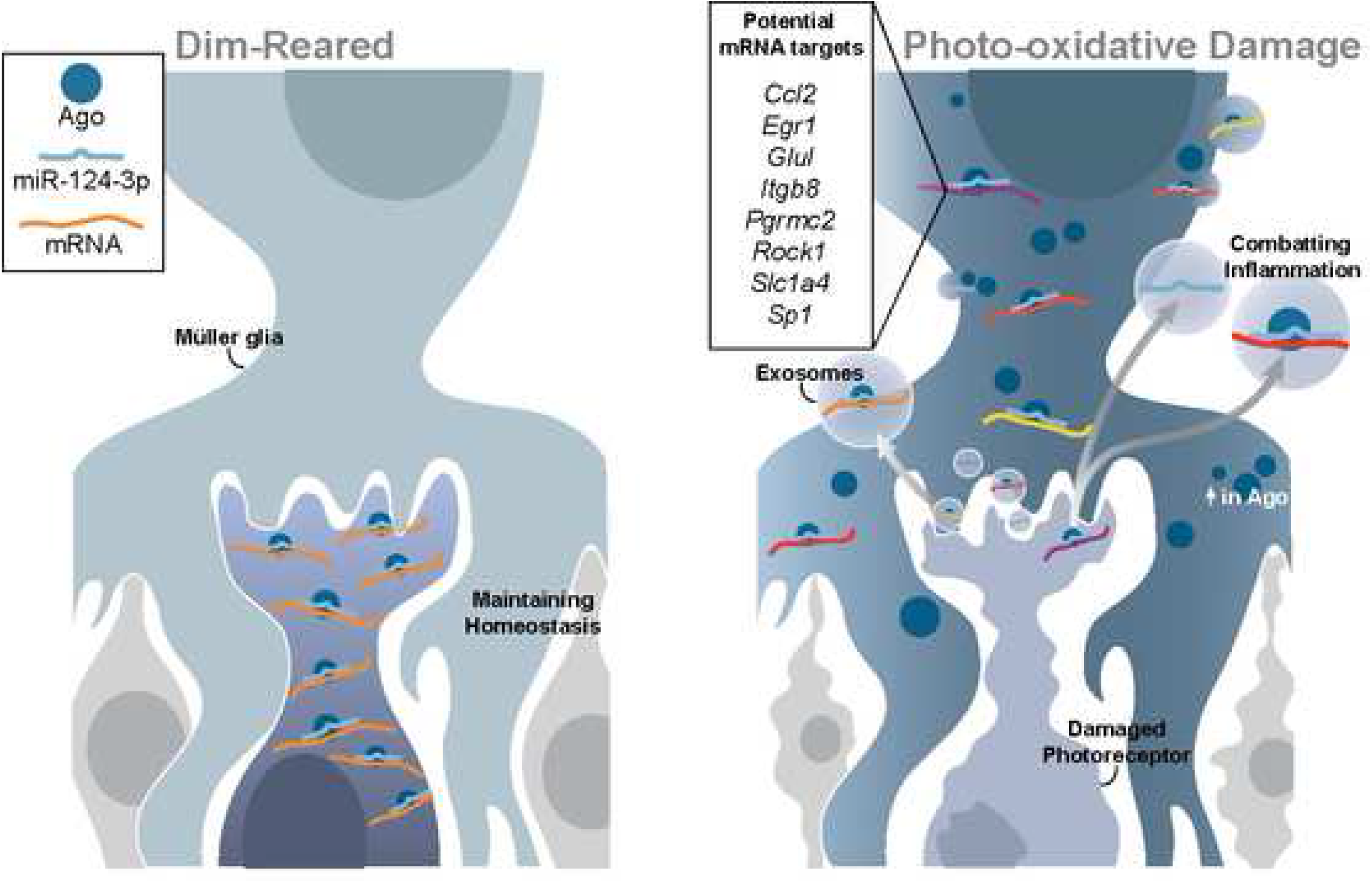
Schematic of retinal miRNA activity in response to PD. Using miR-124-3p as an example, this schematic describes our proposed hypothesis of miRNA activity in response to PD. During DR conditions miRNA maintain homeostasis in the retina, largely in neurons (left panel). In response to PD miRNA are released from the photoreceptors, possibly through exosomes, and translocate to the inner retina (including the Müller glia) to regulate disease-specific processes (right panel). The box highlights potential Müller glia localised mRNA targets (see Figure 6B) of miR-124-3p.

Müller glia are the prominent macroglia in the neural retina. As their processes span across the entirety of the retinal anatomy, they function as homeostatic regulators with an imbalance in the physiology of Müller glia, including ablation of Müller glial miRNA, contributing towards functional alterations and degeneration in the retina (Wohl et al., 2017, Wohl and Reh, 2016, Bringmann and Wiedemann, 2012, Newman and Reichenbach, 1996). Further, it has been shown in the past that miRNA manipulation in Müller glia can determine retinal pathology and even counteract damage (Murray et al., 2013, Zeng et al., 2017). We have shown three separate observations that, taken together may indicate a glial specific miRNA response to inflammation in the retina: 1) enrichment of GO terms and pathways involved in the immune response in the AGO2-bound targetome as aforementioned; 2) upregulation and localisation of AGO2 to the INL where the Müller cell bodies are located; 3) increased binding of miR-124-3p, the most abundant AGO2-bound miRNA, to mRNA targets located in Müller glia following PD. We postulate that as a response to stress, retinal miRNA shift from their usual neuronal maintenance roles in homeostasis, in order to preferentially direct their activity to “Janus-faced” Müller glia that have both beneficial and deleterious features under duress (Bringmann et al., 2009). How this change in miRNA activity occurs is still relatively unknown. However, a number of studies have found that the intercellular transport of miRNA may be mediated by exosomes, small bioactive vesicles ranging from 40-150 nm that are released by cells and play important roles in cell-to-cell communication (Arroyo et al., 2011, Turchinovich et al., 2011, Valadi et al., 2007, Vickers et al., 2011). Further investigations into the activity and translocation of miRNA and more broadly the role that exosomes play in the possible translocations of these dynamic molecules is required to explore the molecular language cells use to communicate and regulate homeostatic and stress responses.

## Conclusion

In conclusion, our data suggests that miRNA are actively involved in regulating key genes and biological pathways leading to retinal degeneration. Whilst the AGO2 HITS-CLIP miRNA gene expression did not change significantly between healthy and damaged retinas we demonstrate that there is a dynamic change in the miRNA target landscape mediated by favouring a different seed binding region. Further, our data provides further support to the notion that perhaps the change in miRNA activity under stress is mediated by a movement of miRNA to the inner layers of the retina. Additional research into this mechanism of movement will help uncover the intricacies of miRNA physiology in neurodegenerative disorders.

## Acknowledgements

This work would not have been possible without the support of the National Health and Medical Research Council of Australia (NHMRC: 1127705), Retina Australia, The Gordon and Gretel Bootes Foundation and The ANU Translational Fellowship.

## Author Contributions

Conceptualization, J.A.C. and R.N.; Methodology, J.A.C, R.N. K.P. and K.H.; Data analysis, J.A.C., Z.F. and H.P.; Investigation, J.A.C., A.V.C. and Y.W.; Writing – Original Draft,

J.A.C. and R.N.; Writing – Review and Editing, J.A.C., R.N. and Y.W.; Supervision, R.N.; Funding Acquisition, R.N, M.R. and J.P.

## Methods

### Animal experiments

All animal experiments were conducted in accordance with the ARVO Statement for Use of Animals in Ophthalmic and Vision Research, and with approval from the ANU Animal Experimentation Ethics Committee (Ethics ID: A2014/56). C57BL/6J mice were used for mice experiments aged between P60-80. Mice were born and reared in individually vented cages (IVCs) in cyclic dim light conditions (12 hours at 5 lux, 12 hours at 0 lux). All animals were age-matched.

### Photo-oxidative damage

Photo-oxidative damage was used to mimic the pathogenesis of atrophic AMD in the animals. The protocol outlined here is based on a previously published paper (Natoli et al., 2016).

Animals were placed into Perspex boxes coated with a reflective interior surface and exposed to 100 K lux of natural white light from light-emitting diodes (LED). Exposure was continuous for a period of 1-7 days with food and water ad libitum. Animals were administered pupil dilator (Minims atropine sulphate 1% w/v; Bausch and Lomb) to both eyes twice a day during the course of the damage paradigm.

### Tissue collection and preparation

Both dim-reared and photo-oxidative damaged animals were culled by CO_2_ exposure. For retinal extraction, the eye is lifted from the socket and, using an ophthalmic knife, cut across the anterior surface of the retina. The lens is removed and the retina extracted using curved forceps. Retinas are immediately immersed in RNAlater (Thermo Fisher Scientific, Waltham, MA) overnight prior to storage at −80°C. Six retinas from three animals were pooled for a single biological sample, n=4 biological replicate samples for both control and damaged retina were obtained. Retinas were diced to a fine suspension in ice-cold PBS and then UV irradiated three times at 400 mJ/cm^2^ (Stratalinker) with the suspension manually swirled between each irradiation to keep cold and to ensure maximal exposed surfaces. Cross-linked tissue was either used directly or flash-frozen and stored at −80°C.

### Ago HITS-CLIP

Argonaute immunoprecipitation (Ago IP) was performed using a protocol adapted from a previous publication (Moore et al., 2014). Buffers, solutions and linker ligation mixes were all made according to (Moore et al., 2014) including manufacturer supplies. Protein A Dynabeads (Thermo Fisher Scientific) were pipetted into RNAse-free 1.5 ml microcentrifuge tubes. Tubes were placed in a magnetic stand to allow the beads to collect on the side of the tube for buffer removal. Beads were washed three times in 1 ml Bead Wash Buffer (BWB). The beads are resuspended in BWB with 50 μg of rabbit anti-mouse IgG bridging antibody (Jackson ImmunoResearch, West Grove, PA). Tubes were rotated end over end at room temperature for 30 minutes and then washed with BWB. This step is repeated substituting IgG for the Anto-pan Ago antibody (clone 2A8; Millipore, Burlington, MA) (4 μl). Antibody-loaded beads were washed three times with 1xPXL ensuring that beads are fully resuspended with each wash.

Cross-linked retinal tissue was suspended with 1xPXL and incubated on ice for 10 minutes before being lysed by gentle mechanical disruption using sterile pestles. 30 μl of RQ1 DNase (Promega, Madison, WI) was added and lysate incubated at 37°C for 5 minutes and rotated at 1000 RPM in a Thermomixer. RNaseA (Affymetrix, Santa Clara, CA) was diluted 1:1000 in 1xPXL, 10 μl added per 1 ml of lysate and incubated at 37°C for 5 minutes. Subsequently lysates are kept ice-cold to minimise further RNase digestion. Lysates were centrifuged on a tabletop centrifuge at 16,000 g for 40 minutes at 4°C. The supernatant was collected, transferred to the antibody-loaded beads and then rotated end over end for 2 hours at 4°C. The supernatant was removed and beads washed in a series of stringent wash steps: three times in 1xPXL; twice in high-salt buffer; twice in high-stringency buffer; 2 times in low-salt buffer; and twice in 1xPNK buffer.

For sequencing all samples were treated with 4 mg/ml of proteinase K (Roche Diagnostics). RNA and small RNA were extracted and enriched in separate fraction from the supernatants using the mirVana kit according to manufacturer’s instructions to separate samples into two groups: small RNA isolated from AGO2:miRNA and AGO2:miRNA:mRNA fractions; and mRNA isolated from AGO2:miRNA:mRNA fractions. RNA concentration was determined using a Qubit 4 Fluorometer (Thermo Fisher Scientific) and RNA quality was measured on a 2100-Bioanalyser using an RNA Nano Assay Chip (Agilent Technologies, Santa Clara, CA). Only samples with an RNA integrity number (RIN) greater than 9.0 were used for sequencing.

To demonstrate the presence of RNA in AGO2 IP complexes gel electrophoresis of 3’ end-labelled AGO2-bound RNA species was performed. For this residual PNK buffer was removed from the AGO2 IP beads before adding desphosphorylation master mix (manufacturer) and resuspending through gentle vortexing. The mixture was incubated at 37°C for 20 minutes, shaking at 1000 RPM for 15 seconds every 2 minutes in a thermomixer. Beads were washed once with 1xPNK buffer, once with 1xPNK+EGTA buffer, then twice with 1xPNK buffer. 3’ linker ligation master mix was added to the beads and incubated at etc. Beads were washed with high-salt buffer, twice with 1xPNK buffer and then three times with 1xPNK+EGTA buffer. Residual buffer was removed, beads resuspended in 1xLDS sample-loading buffer (Thermo Fisher Scientific) and incubated at 70°C for 15 minutes, shaking at 1000 RPM in a Thermomixer. Beads were collected by centrifugation and the supernatant loaded onto a Novex NuPAGE Bis-Tris 8% gel (Invitrogen, Carlsbad, CA). Proteins were separated at 175 V for 3-4 hours at 4°C and then transferred to nitrocellulose membrane (BioRad) by using a Criterion blot cell (BioRad) for 1 hour at 90 V in 1 x NuPAGE transfer buffer (Invitrogen) containing 10% methanol. The Nitrocellulose membrane was rinsed in phosphate buffered saline and placed in a phosphorimage cassette and a film (GE Healthcare, Chicago, IL) exposed for 24 hours. The film was imaged using a Phosphorimager (Typhoon FLA 9500, GE Healthcare).

### Library preparation, high throughput sequencing and read processing

Library preparation and RNA sequencing was performed by the John Curtin School of Medical Research Biomolecular Research Facility. The libraries were prepared using the CATS Small RNA-seq Kit and CATS RNA-seq Kit according to manufacturer’s instructions (Diagenode, Denville, NJ) for miRNA and mRNA populations, respectively. Libraries were sequenced on an Illumina HiSeq 2500 acquiring single-end read lengths of 51 bp for the global miRNA and HITS-CLIP experiments and 76 bp for the global mRNA experiment. Raw reads were processed with cutadapt (settings) to trim adaptors, artefact bases and barcodes. FASTQC sequence quality analysis were performed before and after cleaning. The average clean read length was 20 nt and 45 nt for miRNA and mRNA libraries, respectively. All raw sequencing data have been deposited onto the Sequence Read Archive (PRJNA606092) as part of the National Center for Biotechnology Information and will be released upon publication.

### Global mRNA read mapping

Clean reads were aligned to mm10 genome with bwa-mem (Li and Durbin, 2010) and samtools (Li et al., 2009) used for sorting and statistical analysis. Mapped reads were assigned to annotated mm10 RefSeq genes based on FeatureCounts (Liao et al., 2014). Trimmed Mean of M-values (TMM) (Law et al., 2014, Ritchie et al., 2015, Robinson and Oshlack, 2010) was used for normalisation. Differential expression was analysed using the voom-limma package (Ritchie et al., 2015) with a linear model fit and Bayes’ adjustment of p-values.

### Global miRNA read mapping

Clean reads were aligned using bwa aln to the mouse mature miRNA sequences in miRBase (Kozomara and Griffiths-Jones, 2014, Li and Durbin, 2010). Alignment was summarised with Unix shell script (awk and grep). The miRNAs were ranked based on their mean expression level and normalised by library size. Top ranking miRNA targets were identified using TargetScan (Lewis et al., 2005) and miRNet (Fan et al., 2016) and cross-referenced with out AGO:mRNA data set. For a robust differential expression analysis only miRNAs with CPM>10 in at least four out of the eight samples were used. Voom-limma package was used with sample quality weights, Trimmed Mean of M-values (TMM) (Law et al., 2014, Ritchie et al., 2015, Robinson and Oshlack, 2010) used for normalisation.

### Enrichment analyses

GSEA (Gene Set Enrichment Analysis) was performed with camera (Wu and Smyth, 2012) against the gene sets listed in the Hallmark and C2 pathways from the Molecular Database (MsigDB v5) and GO database. GOrilla (Eden et al., 2009) was also used to generate enrichment scores for pathways and GO terms. Terms with >3 genes and P<0.05 were deemed significantly enriched. For identification of potentially regulated miRNA for the differentially expressed genes, the genes were input into miRNet database and cross-referenced with our miRNA global analysis.

### Analysis of HITS-CLIP read mapping

Raw reads were pre-processed with similar methods as the global mRNA and global miRNA sequence outputs. Clean reads were aligned using bwa aln to mouse mature miRNA and pri-miRNA sequences downloaded from miRbase and the RNA sequences downloaded from the mm10 genome, respectively (Li and Durbin, 2010). awk, samtools (Li et al., 2009) and FeatureCounts (Liao et al., 2014) were used for count sorting and summary. The mean mRNA and miRNA counts were ranked normalized CPM and only highly expressed (conditions) miRNAs or mRNAs used for differential expression and pathway analyses.

### Peak calling and sequence motif analysis of miRNA binding regions

The mm10 genome alignment files were converted to the bed format and binding peaks were called with Homer (findPeaks with factor and tss styles, http://homer.ucsd.edu/homer/), separately, for PD group and DR group. The peaks were annotated using Homer (annotatePeaks.pl), extracted and separated into different regions (exon, intron, transcription termination site (TTS), ncRNA, miRNA, 5’ UTR, 3’ UTR). Then the enriched sequence motifs were searched within the sequences of the 3’ UTR with Homer (findMotifs.pl). The significantly expressed miRNAs and their targets were matched based on the motifs, their expression levels, and cross-referenced with our results from global miRNA and mRNA experiments.

### Western Blot

Retinas from dim-reared (DR) mice (n=6) and mice exposed to 5 days of photo-oxidative damage (PD) (n=6) were lysed in CellLytic^TM^ Cell Lysis Buffer (Sigma-Aldrich) supplemented with a protease inhibitor cocktail (Sigma-Aldrich). A total of 20 μg of protein was denatured in LDS loading buffer (Thermo Fisher Scientific) and subjected to electrophoresis on a Novex™ 4-20% Tris-Glycine Mini Gel (Thermo Fisher Scientific). The protein bands were transferred to a nitrocellulose membrane (Bio-Rad, CA, USA), blocked in 3% BSA in PBS for 1 hours and then incubated with primary antibody (anti-AGO2 antibody (1:1000): ab32381, Abcam, Cambridge, UK; anti-GAPDH antibody (1:5000): G9545, Sigma-Aldrich) overnight at 4°C. Membranes were washed in PBS/0.1 % Tween (Sigma-Aldrich and then incubated with HRP-conjugated Goat Anti-Rabbit IgG (H+L) antibody for 2 hours. The signal was developed with the ClarityTM Western ECL Substrate (Bio-Rad) and imaged using the ChemiDoc^TM^ MP Imaging System with Image Lab^TM^ software (Bio-Rad). The intensity of the AGO2 bands was first normalised to GAPDH and then expressed as a ratio of dim-reared controls.

### Immunohistochemistry

Immunohistochemistry was performed as previously published (Chu-Tan et al., 2018). After enucleation, the eyeballs were fixed in 4% paraformaldehyde (Sigma-Aldrich), dehydrated in 15% sucrose (Sigma-Aldrich) and embedded in Tissue-Tek® Optimal Cutting Temperature Compound (Sakura Finetek, California, USA) in a consistent orientation. Cryo-sectioning was carried out on a Leica CM 1850 cryostat at 12 μm thickness and the tissue sections were mounted on SuperFrost Ultra Plus^TM^ Adhesion Slides (Thermo Scientific) and stored at − 20°C until use.

When required for immunohistochemistry, the slides were first rehydrated in Milli-Q water then equilibrated in PBS. Antigen retrieval was performed with 100% Revealit-Ag Antigen Recovery Solutions (ImmunoSolutions, QLD, Australia) at 37°C for 45 minutes then the crysections were blocked in 10% Normal Goat Serum (Sigma-Aldrich, MO, USA). The cryosections were then incubated in anti-AGO2 antibody (1:1000, ab32381, Abcam, Cambridge, UK) overnight at 4°C. The next day, the cryosections were incubated with Alexa Fluor^TM^ 488 goat-anti-Rabbit IgG (1:1000) for 2 hours at room temperature and then the nuclei counterstained with Hoechst 33342 (Sigma-Aldrich). The cryosections were imaged using the A1+ Confocal Microscope System with NIS-Elements Advanced Research software (Nikon, Tokyo, Japan). For each cryosection, 1 μm thick z-stacked images were captured, the final image was obtained by compressing all focal planes using the maximum intensity projection processing function in ImageJ (NIH, USA).

### cDNA preparation and quantitative real-time PCR

Complementary DNA (cDNA) used for in *situ* hybridisation and quantitative PCR (qPCR) was synthesised using the Tetro cDNA synthesis kit (Bioline, London, UK) with oligo-dT primers according to the manufacturers’ instructions.

qPCR was performed using an AGO2 TaqMan hydrolysis probe (Mm00838341_m1, ThermoFisher Scientific, MA, USA) as previously published (Chu-Tan et al., 2018). *Gapdh* (Mm99999915_g1, ThermoFisher Scientific) was used as a reference gene.

### In situ hybridisation

A 445 bp region of the mouse *AGO2* mRNA (NM_153178.4) was amplified from retinal cDNA using the forward primer GAGACAGTCCACCTCTTGTGG and reverse primer GCCCAGAAGCAAACAACACC. The reverse primer was extended with a T7 RNA Polymerase tag ATATATTAATACGACTCACTATAGG at the 5’ end. A total of 35 PCR cycles [melting (95°C, 15 seconds), annealing (60°C, 15 seconds), extension (72°C, 10 seconds)] were carried out and the presence of the correct amplicon was verified using gel electrophoresis. After amplification, the PCR product was mixed with ¼ of ammonium acetate and 10 volumes of ice-cold absolute ethanol. The PCR product centrifuged (13000g, 15 mins, 4°C), then the pellet was washed with 70% ice-cold ethanol and resuspended (13000 g, 2 mins, 4°C). The ethanol was removed and the PCR product was reconstituted in Ultrapure water (Gibco, Thermo Fisher Scientific, MA, USA). The purity and concentration of the reconstituted PCR product was assessed on a ND-1000 spectrophotometer (Nanodrop Technologies, DE, USA). A riboprobe was transcribed from the template using T7 RNA polymerase (Promega, Madison, WI) incorporating digoxigenin (DIG) (SP6/T7; Roche, Basel, Switzerland), as previously published (Cornish et al., 2005). In situ hybridisation was performed on cryosectioned retinas using previously described protocols (Cornish et al., 2005). Hybridisation of the AGO2 probe was achieved at 59°C overnight, unbound probe removed by washing with 20 x saline sodium citrate (pH 7.4) at 60°C and developed using nitro blue tetrazolium and 5-bromo-4-chloro-3-indolyl phosphate (NBT/BCIP) (Sigma Aldrich) for 60 minutes. The reaction was stopped by washing with ultrapure water and the slides were mounted with Aqua-Poly/Mount.

## Supplemental Tables

**Supplementary Table 1.** C2 Gene Set Enrichment Analysis on differentially expressed AGO2-bound mRNA between DR and PD retinas.

| Pathway | Up or Down | Number of genes | P-value |
| --- | --- | --- | --- |
| Rhodopsin Pathway | Down | 17 | 7.20e-10 |
| Interferon Alpha Beta Signaling | Up | 17 | 2.40e-4 |
| EGF Signaling | Up | 38 | 2.90e-5 |
| Opsins | Down | 6 | 2.70e-4 |
| Initial Triggering of Complement | Up | 5 | 3.20e-4 |
| Classic Pathway | Up | 6 | 4.00e-4 |
| Complement Pathway | Up | 6 | 4.00e-4 |
| Regulation of complement cascade | Up | 8 | 2.10e-3 |
| NF-KB Pathway | Up | 19 | 3.90e-3 |
| Stress Pathway | Up | 16 | 6.20e-3 |
| Tumour Necrosis Factor Alpha | Up | 102 | 3.45e-3 |
| Inflammatory Response to LPS | Up | 25 | 1.50e-2 |

**Supplementary Table 2.** AGO2-bound mRNA predicted to bind to miR-124-3p.

| Gene | logFold Change in PD vs DR | P-value |
| --- | --- | --- |
| *Antxr2* | 2.3626328 | 9.70E-09 |
| *Myo10* | 1.798979023 | 1.95E-08 |
| *Stk35* | -0.939450988 | 3.55E-05 |
| *Ammecr1* | -0.994602386 | 4.59E-05 |
| *Plekhf2* | -0.876329158 | 5.98E-05 |
| *Slc16a1* | -0.829380386 | 7.58E-05 |
| *Egr1* | 1.90922451 | 0.000124661 |
| *Zcchc24* | 0.914702034 | 0.000136088 |
| *Slc1a4* | 1.160994865 | 0.000177268 |
| *Tor3a* | 1.363762931 | 0.00027897 |
| *Slc2a13* | 0.891574534 | 0.000858168 |
| *Acaa2* | 1.114278205 | 0.000860691 |
| *Fam199x* | -0.506414329 | 0.002787212 |
| *Tob2* | -0.497194078 | 0.002837396 |
| *Ccl2* | 2.711785452 | 0.00352564 |
| *Glul* | -0.408204948 | 0.004504003 |
| *Thrb* | -2.493894838 | 0.007497924 |
| *Fam53b* | -2.113053204 | 0.007877904 |
| *Ccdc117* | -0.505556285 | 0.009372384 |
| *Anxa5* | 0.79985392 | 0.011338264 |
| *Itgb8* | 1.124728236 | 0.011993412 |
| *Fam89a* | -0.797404686 | 0.012763135 |
| *Xkr4* | 0.638058956 | 0.013658401 |
| *Ascc2* | 0.683486851 | 0.01366163 |
| *Klf6* | 0.624593326 | 0.01662975 |
| *Lonrf1* | 0.444487282 | 0.016767637 |
| *Nbeal2* | 0.795824543 | 0.017171796 |
| *Pgrmc2* | 0.440415689 | 0.018366563 |
| *Tiprl* | -2.140248833 | 0.019541074 |
| *Rgs9* | -0.33650089 | 0.021885195 |
| *Klhl28* | -0.614793382 | 0.025672891 |
| *Slc7a8* | 0.687754313 | 0.026484867 |
| *Pik3ca* | -1.808157219 | 0.026708444 |
| *Tmcc3* | 1.986152996 | 0.029708159 |
| *Spry3* | -0.501623413 | 0.029870207 |
| *Nr1d2* | -0.354795692 | 0.030021147 |
| *Rnf11* | -0.365613023 | 0.03078371 |
| *Lrrc58* | 0.219233176 | 0.031143983 |
| *Sigmar1* | -0.624054103 | 0.033853882 |
| *Rock2* | 0.882579372 | 0.035479855 |
| *Ncoa2* | -2.043173774 | 0.036152073 |
| *Srsf6* | 0.482334496 | 0.039718188 |
| *Rock1* | -2.158513569 | 0.039728568 |
| *Mmp16* | 0.973148135 | 0.039863984 |
| *B4galt6* | 0.387926472 | 0.040073129 |
| *Rbm20* | -2.00730089 | 0.040365883 |
| *Myh9* | 0.288650483 | 0.0407971 |
| *Sptlc2* | 0.37871085 | 0.040926428 |
| *Dusp6* | 0.944425222 | 0.041557607 |
| *Nr3c1* | -1.757589789 | 0.042048432 |
| *Ttc9* | 0.420767978 | 0.04213689 |
| *Mapre1* | 0.329427138 | 0.042708551 |
| *Arih1* | 0.715195672 | 0.044913444 |
| *Oxsr1* | -2.063965828 | 0.045342592 |
| *Sp1* | 1.834869082 | 0.045361948 |
| *Jag1* | -1.171294072 | 0.045984879 |
| *Ncor2* | 1.879010576 | 0.047358544 |
| *Tub* | -0.30229807 | 0.047541267 |
| *Sertad3* | 0.815671744 | 0.048025558 |
| *Stt3a* | -0.342667898 | 0.048466343 |
| *Csgalnact2* | 0.942186647 | 0.048469156 |

